# Estimating contemporary migration numbers of adults based on kinship relationships in iteroparous species

**DOI:** 10.1101/2022.03.09.483709

**Authors:** Tetsuya Akita

## Abstract

In this paper, we describe the development of estimators for the contemporary migration number and rate of adults between two populations in iteroparous species. The proposed estimators are based on known half-sibling (HS) and/or parent–offspring (PO) relationships observed between the populations across breeding seasons. The rationale is that the HS and PO pairs exhibit information about the occurrence frequency of parental movements during the breeding interval. The proposed method allows for variance in the average number of offspring per parent both within and between populations. In addition, coupled with the PO pairs found within the population, the estimators can be obtained from only genetic data. Generally, a sample size representing the square root of the population size is required to obtain meaningful information regarding migration. We describe a detailed evaluation of the performance of the estimators by running an individual-based model, and the results provide guidance regarding sample sizes to ensure the required accuracy and precision. Furthermore, given that there are few effective methods to estimate adult movement (especially when populations cannot be genetically distinct), we discuss the usefulness of this proposed kinship assignment method in terms of conservation biology and wildlife management.

## 1 INTRODUCTION

In conservation and wildlife management, estimating current ecological information is essential for monitoring population levels and proposing effective management strategies. The contemporary migration rate, i.e., the recent rate of movement of individuals or gametes between predefined populations, is a critical component when determining the current degree of gene flow and synchrony among the population (Lowe & Allendorf, 2010; Waples & Gaggiotti, 2006). Therefore, estimating the contemporary migration rate provides information about the degree of genetic differentiation and demographic dependency, which can delineate evolutionarily significant units and identify appropriate management units (Moritz, 1994; Palsbøll, Bérubé, & Allendorf, 2007). In addition, the availability of migration rate can link to underlying population dynamics, allowing for more precise and flexible evaluations of management practices, e.g., stock assessment in fisheries (Hampton & Fournier, 2001) and invasive species control (Sakai et al., 2001).

Essentially, there are two approaches to estimate the contemporary migration rate. The first approach involves using the mark-recapture (MR) method, which estimates the migration rate along with other population parameters (Kéry & Schaub, 2011; Thorson et al., 2021). Here, the rational is that, for example, conventional tags provide release and recovery location information for known release and recovery dates, which generates a movement fraction matrix among strata per given time interval. This direct method is relatively easy to interpret; however, it is generally hampered by several uncertainties, e.g., tag loss, tagging-related mortality, and time-varying reporting rates (Hilborn & Walters, 1992). In addition, the MR method is generally limited in practice to situations where collection of a sufficient number of adult samples at multiple sites is possible over a short period.

The second approach involves using numerous genetic markers to assign individuals to source populations, thereby allowing the inference of recent migration (since Paetkau, Calvert, Stirling, & Strobeck, 1995). The previously developed model of Wilson and Rannala (2003), which is one of the population assignment methods and implemented in a software ‘BayesAss’, can output the point estimate as an element of the current migration matrix. However, according to several papers (Faubet, Waples, & Gaggiotti, 2007; Wang, 2014), the accuracy of the migration rates estimated by this model is valid only when populations are highly differentiated (e.g., *F*_ST_ ≥ 0.05), which suggests that it may be difficult to infer the degree of demographic dependency among populations because demographic independency is realized even if there are many migrants.

Parental assignments, which are based on genetic markers and are frequently used to complement the population assignment method in ecological studies, also provide information about the current level of migration. In contrast to the population assignment method, the parental assignment method does not require population differentiation (Wang, 2014); however, the current methodology focuses on estimating dispersal kernels and, to estimate the migration rate, requires the assumption that adults are not migrant, e.g., pollen/seed dispersal in plants (Ashley, 2010) or larval drift in marine animals (Gagnaire et al., 2015). Thus, this method may not consider the adult movement of iteroparous species, i.e., multiple reproductive cycles during the lifetime, which may change the spawning ground for each breeding season.

Close-kin mark-recapture (CKMR) is a recently developed method to estimate adult population size along with life-history parameters that utilizes the known kinship information in a sample (Waples & Feutry, in press, and references therein). In the CKMR method, the presence of a kinship pair in the sample is analogous to the recapture of a marked individual in the MR method. Note that kinship pairs in the sample are less likely to be observed in larger populations; thus, the number of kinship pairs may reflect the number of adults in the population (Bravington, Skaug, & Anderson, 2016). While recent studies of CKMR argued that availability of kinship information to identify metapopulation structure (Conn, Bravington, Baylis, & Ver Hoef, 2020; Trenkel, Charrier, Lorance, & Bravington, 2022), CKMR-based migration estimation has not been well developed or examined extensively in a simulation study.

Thus, in this paper, we propose a method to estimate the contemporary migration number or rate of adults between two predefined populations in iteroparous species, where the migration direction is specified. Under the assumption that kinships are genetically detected without error, the proposed method is based on the numbers of half-sibling (HS) and parent–offspring (PO) pairs in a sample. Here, sampling can be either invasive or noninvasive and is completed at two breeding seasons; sampling offspring (young-of-year individuals) and parents likely share a PO relationship with the offspring sampled in one population at the first breeding season and in the other population at the second breeding season. The rationale for estimating the migration number or rate is that the number of HS and PO pairs found between populations contains information about the frequency of parental movements during the breeding interval. Our model explicitly incorporates reproductive variation within and between populations, thereby making it possible to target a species whose fertility is affected by environmental differences between populations. First, we explain the modeling assumption and sampling scheme. Then, we analytically determine the estimators of the contemporary migration number or rate of adults, which are based on the numbers of HS and/or PO pairs. Finally, by running an individual-based model, we investigate the performance of the estimator and provide a guide for a sample size. Note that the proposed modeling framework can be applied to diverse animal species; however, the description of the model focuses on fish species, which are presently the best target candidate for the proposed method.

## 2 Theory

Here, we present the theoretical foundation of estimating a migrant number and migration rate of iteroparous species using HS or PO pairs found between two populations with different sample timing. Note that the estimators can hold under flexible assumptions for reproduction (**Appendix 2**); however, here, we assume a relatively simple case. The main symbols used in this paper are summarized in **Table 1**.

**Table 1:**
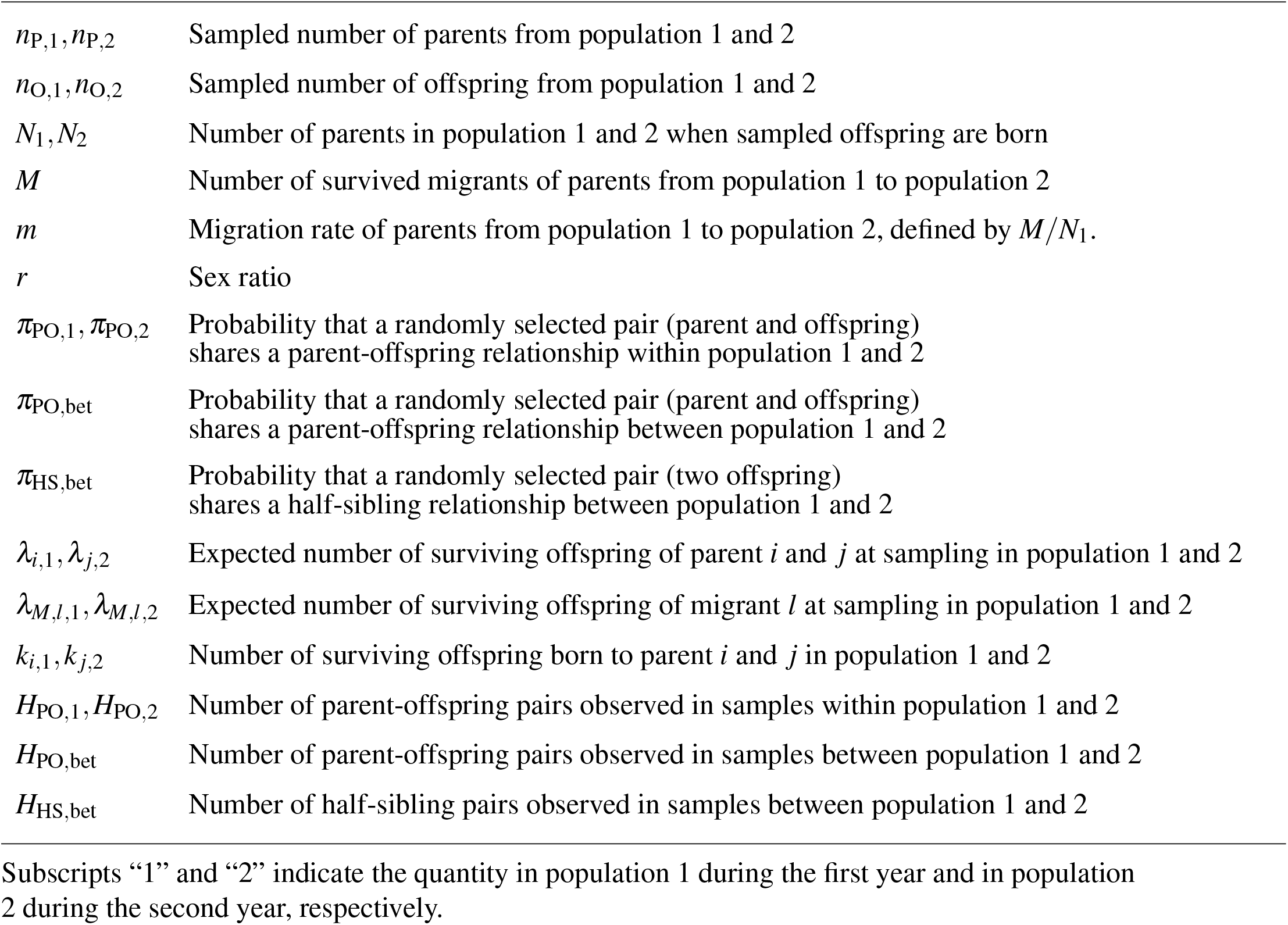
The list of mathematical symbols employed in the main text

|  |  |
| --- | --- |
| $n_{P,1}, n_{P,2}$ | Sampled number of parents from population 1 and 2 |
| $n_{O,1}, n_{O,2}$ | Sampled number of offspring from population 1 and 2 |
| $N_1, N_2$ | Number of parents in population 1 and 2 when sampled offspring are born |
| $M$ | Number of survived migrants of parents from population 1 to population 2 |
| $m$ | Migration rate of parents from population 1 to population 2, defined by $M/N_1$. |
| $r$ | Sex ratio |
| $\pi_{PO,1}, \pi_{PO,2}$ | Probability that a randomly selected pair (parent and offspring) shares a parent-offspring relationship within population 1 and 2 |
| $\pi_{PO,bet}$ | Probability that a randomly selected pair (parent and offspring) shares a parent-offspring relationship between population 1 and 2 |
| $\pi_{HS,bet}$ | Probability that a randomly selected pair (two offspring) shares a half-sibling relationship between population 1 and 2 |
| $\lambda_{i,1}, \lambda_{j,2}$ | Expected number of surviving offspring of parent $i$ and $j$ at sampling in population 1 and 2 |
| $\lambda_{M,l,1}, \lambda_{M,l,2}$ | Expected number of surviving offspring of migrant $l$ at sampling in population 1 and 2 |
| $k_{i,1}, k_{j,2}$ | Number of surviving offspring born to parent $i$ and $j$ in population 1 and 2 |
| $H_{PO,1}, H_{PO,2}$ | Number of parent-offspring pairs observed in samples within population 1 and 2 |
| $H_{PO,bet}$ | Number of parent-offspring pairs observed in samples between population 1 and 2 |
| $H_{HS,bet}$ | Number of half-sibling pairs observed in samples between population 1 and 2 |

Subscripts “1” and “2” indicate the quantity in population 1 during the first year and in population 2 during the second year, respectively.

### 2.1 Hypothetical population

Assume that we have a set of populations 1 and 2, where random mating occurs within each population and parents can move to the other population after the reproductive season ends. Without loss of generality, we consider the movement of parents from population 1 to population 2, and we focus on estimating the migration number or rate. In this framework, we demonstrate that considering two reproductive seasons (hereafter referred to as “the first year” and “the second year”) is sufficient to estimate the migration number or rate. **Figure 1a** shows a schematic representation of the kinship relationships and parent movements.

**FIGURE 1.**
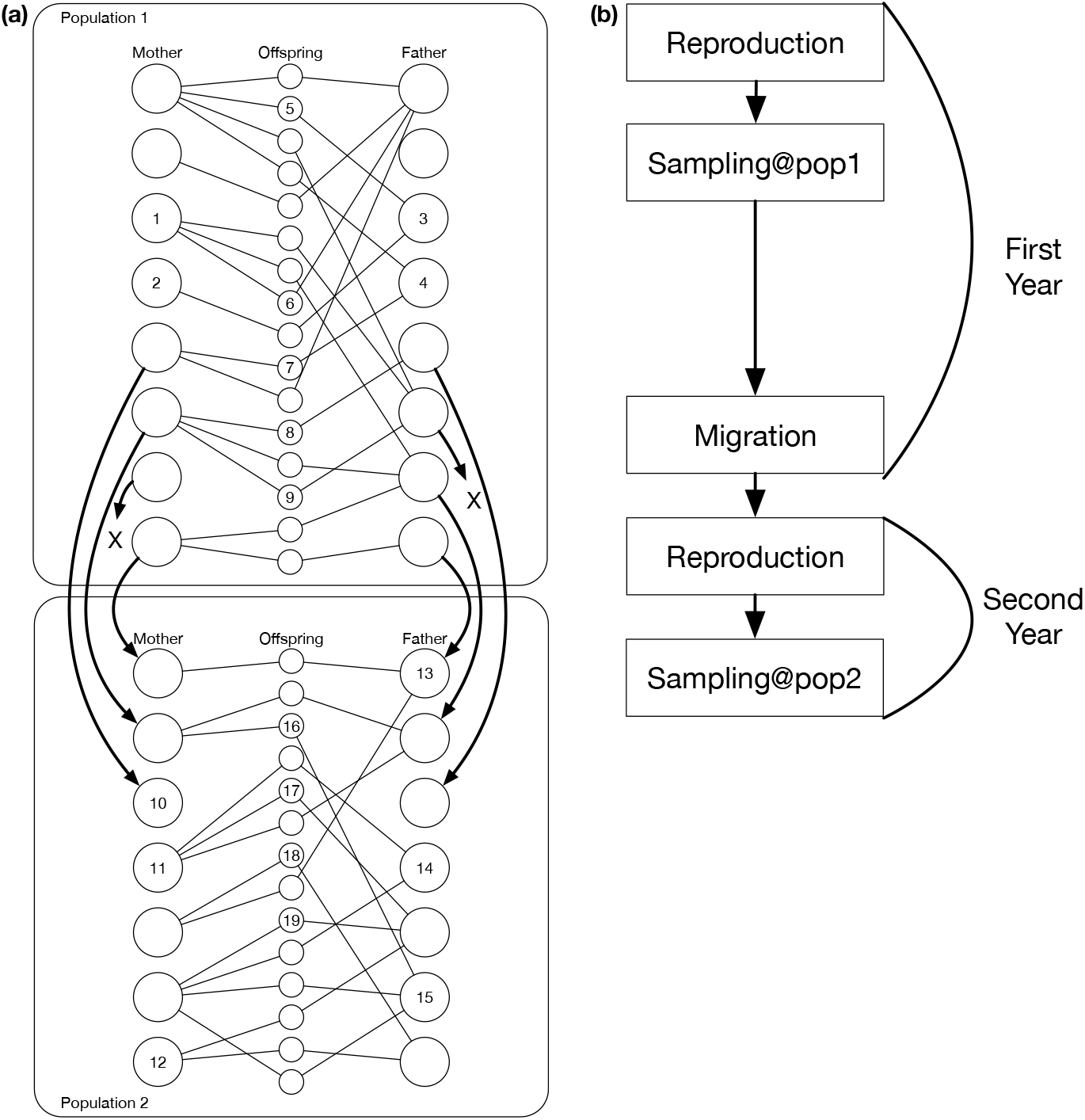
(a) Hypothetical populations with *N*_1_ = 16, *N*_2_ = 14, and *M* = 6. Upper and lower areas indicate individuals in population 1 before migration (at the sample timing in the first year) and individuals in population 2 after reproduction (at the sample timing in the second year), respectively. Open circles on the left, right, and center represent mothers, fathers, and their offspring, respectively. The thin line denotes PO relationship. Bold arrows denote migration, and x denotes failure to survive at the sampling in the second year. Sampled individuals are labeled with an index number. Number of sampled individuals in this example: *n*_P,1_ = 4, *n*_O,1_ = 5, *n*_P,2_ = 6, *n*_O,2_ = 4; numbers of kinship pairs: *H*_PO,bet_ = 1 (i.e., “7-10” pair), *H*_HS,bet_ = 2 (i.e., “8-16” and “9-16” pairs), *H*_PO,1_ = 3 (i.e., “1-6”, “3-5” and “4-7” pairs), and *H*_PO,2_ = 2 (i.e., “11-17” and “15-16” pairs). (b) Phases of events relevant to this study in the given timeline.

Assume that there are *N*_1_ parents in population 1 at the beginning of the first year. Each parent produces a number of offspring that is governed by the parent’s reproductive potential, which is denoted by *λ*_*i*,1_ (*i* = 1, 2, …, *N*_1_). After the reproductive season, some parents begin to move toward population 2. Here *M* survived migrants arrive in population 2; thus, there are *N*_2_ parents in population 2 at the beginning of the second year (*M* ≤ *N*_1_ and *M* ≤ *N*_2_). Similar to population 1, *N*_2_ parents produce a number of offspring governed by *λ* _*j*,2_ (*j* = 1, 2, …, *N*_2_). Note that reproductive potential is determined by several factors. Additional details about reproductive potential can be found in **Appendix 1**. Theoretically, the number of offspring for each parent is set as a random variable with mean *λ*_*i*,1_ or *λ* _*j*,2_.

### 2.2 Sampling

To estimate the migration number or rate, we utilize the number of HS and/or PO pairs observed in a sample. In both populations, *n*_O,1_ and *n*_O,2_ offspring are sampled randomly in the first and second year, respectively, which are identified as young-of-year individuals without error. In addtion, *n*_P,1_ and *n*_P,2_ parents are sampled randomly immediately after the end of the reproductive season in the first and second years, respectively. For mathematical tractability, the parents must survive the reproductive season; therefore, both the mother and father of a give offspring have the potential to be sampled in the same year. Note that all the four sample types, i.e., *n*_P,1_, *n*_P,2_, *n*_O,1_, and *n*_O,2_, are not always required to estimate the migration number or rate. The types of required samples depends on the given situation, which is explained later. Up to subsection 2.6, we focus on only the HS and PO pairs found between populations 1 and 2. To apply the proposed method to the fishery assessment, we assume an invasive sampling procedure, which prevents us from finding a PO pair such that a parent and offspring is sampled in population 1 and 2, respectively. **Figure 1b** shows the timeline of the sampling scheme for these hypothetical populations.

In the example shown in **Figure 1a**, five offspring and four parents are sampled in population 1, four offspring and six parents are sampled in population 2, and two HS pairs and a single PO pair are observed between the two populations 1 and 2. In addition, there are several PO pairs observed within the same population, which are available to estimate parent numbers, which is explained in subsection 2.7. In our modeling framework, if a full-sibling relationship is found, we count it as two HS pairs. The numbers of HS and PO pairs found between the two populations are determined via pairwise comparison of all the sample individuals, i.e., comparisons of *n*_O,1_ × *n*_O,2_ and *n*_O,1_ × *n*_P,2_, respectively.

### 2.3 HS pair-based model

Here, we consider the probability that two offspring sampled in populations 1 and 2 will share a HS relationship with an arbitrary mother or father, denoted by π_HS,bet_. Note that we assume equal reproductive potential among parents (up to subsection 2.7) although this assumption can be relaxed for most of the case (see **Appendix 2** for additional information). Here, π_HS,bet_ can be partitioned into three probabilities, i.e., (i) the probability that sampled offspring in population 1 born to a parent that safely arrives in population 2 (hereafter, referred to as a “migrant”); (ii) the probability that sampled offspring in population 2 are born to a migrant; and (iii) the probability that the migrant of a sampled offspring in population 1 and the migrant of a sampled offspring in population 2 are identical.

To assess these probabilities, recall that there must be two parents of arbitrary offspring. The first probability is the sum of the father–offspring and mother–offspring relationship, which can be expressed as *r*_M_*M/*(*r*_S,1_*N*_1_) +(1 − *r*_M_)*M/*((1 − *r*_S,1_)*N*_1_), where *r*_M_ and *r*_S,1_ indicate the sex ratio of the parents in the migrants and in all of the population 1, respectively. Under the assumption that *r*_M_ = *r*_S,1_, the probability is simplified to 2*M/N*_1_. Similarly, the second probability is simplified to 2*M/N*_2_, which implies the assumption regarding an equal sex ratio in migrants and others. The third probability corresponds to the probability that two randomly selected migrants are identical, i.e., 1*/M*. Taken together, we obtain the following:

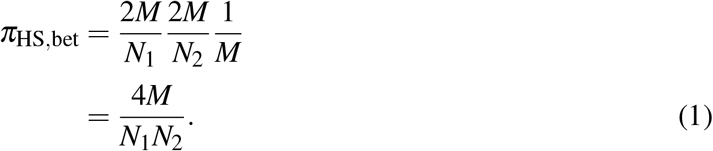

This form holds under a flexible setting for the reproductive potential, e.g., a situation with random variable of *λ*_*i*,1_ and *λ* _*j*,2_ (**Appendix 2**). If all *N*_1_ parents safely move to population 2 (i.e., *M* = *N*_1_), π_HS,bet_ equals 4*/N*_2_. In addition, if population 2 comprises only migrants (i.e., *M* = *N*_2_), π_HS,bet_ equals 4*/N*_1_. In extreme cases, these probabilities take similar forms as the HS probability, which is sampled randomly from two different cohorts within a population (Bravington, Skaug, & Anderson, 2016).

Let *H*_HS,bet_ be the number of HS pairs found in the offspring samples of size *n*_O,1_ and *n*_O,2_. Assuming the total number of HS pairs between the two populations much greater than *H*_HS,bet_, the distribution is approximated by a binomial form (i.e., *H*_HS,bet_ ∼ Binom[*π*_HS,bet_, *n*_O,1_*n*_O,2_]). Thus, the theoretical expectation of *H*_HS,bet_ is given as follows:

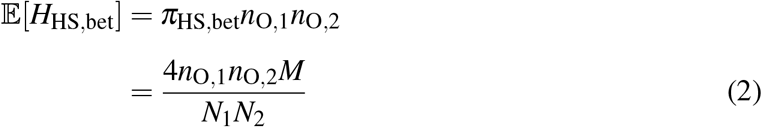

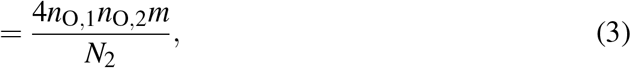

where *m* is the migration rate satisfying *M* = *mN*_1_. The observed number of HS pairs in a sample found between the populations 1 and 2 is defined by 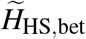, and 𝔼[*H*_HS,bet_] in Equation 2 is replaced by 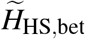, thereby generating the linear estimator of *M*:

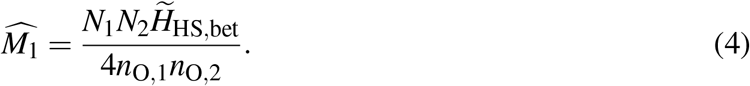

In this paper, a “tilde” and “hat” symbols indicate the observation and estimator of a variable, respectively. Similarly, 𝔼[*H*_HS,bet_] in Equation 3 is replaced by 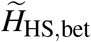, which generates the linear estimator of *m*:

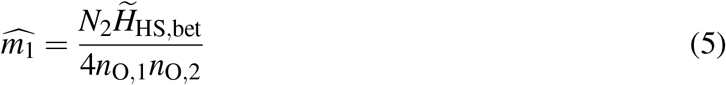

The subscripts associated with these estimators indicate the numbering of the proposed estimators, which is summarized in Table 2.

**Table 2:** Summary of proposed estimators for required parameters and kinship types

| Estimator | $N_1$ | $N_2$ | $n_{O,1}$ | $n_{O,2}$ | $n_{P,1}$ | $n_{P,2}$ | Required kinship type |
| --- | --- | --- | --- | --- | --- | --- | --- |
| $\widehat{M}_1$ | given | given | ✓ | ✓ | | | HSP |
| $\widehat{M}_2$ | given | given | ✓ | | | ✓ | POP |
| $\widehat{M}_3$ | given | given | ✓ | ✓ | | ✓ | HSP & POP |
| $\widehat{M}_4$ | given | estimated | ✓ | ✓ | | ✓ | HSP & POP |
| $\widehat{M}_5$ | estimated | estimated | ✓ | ✓ | ✓ | ✓ | HSP & POP |
| $\widehat{m}_1$ | — | given | ✓ | ✓ | | | HSP |
| $\widehat{m}_2$ | — | given | ✓ | | | ✓ | POP |
| $\widehat{m}_3$ | — | given | ✓ | ✓ | | ✓ | HSP & POP |
| $\widehat{m}_4$ | — | estimated | ✓ | ✓ | | ✓ | HSP & POP |

HSP: half-sibling pair; POP: parent–offspring pair.

### 2.4 PO pair-based model

In the folowing, we consider the probability that offspring sampled in population 1 and a parent sampled in population 2 share a PO relationship, denoted by π_PO,bet_. π_PO,bet_ can be partitioned into the above mentioned probabilities. Note that probabilities (i) and (iii) are the same as that introduced in the previous subsection, i.e., 2*M/N*_1_ and 1*/M*. In addition, probability (ii) is *M/N*_2_ by definition. Taken together, we obtain the following:

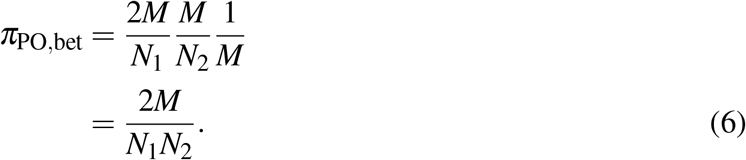

This form also holds under a flexible setting for the reproductive potential, as noted in the **Appendix 2**. Here, if all *N*_1_ parents safely move to population 2 (i.e., *M* = *N*_1_), *π*_PO,bet_ equals 2*/N*_2_. In addition, if population 2 comprises only migrants (i.e., *M* = *N*_2_), *π*_PO,bet_ equals 2*/N*_1_. In extreme cases, these probabilities take similar forms to the PO probability that is sampled randomly within a population (Bravington, Skaug, & Anderson, 2016).

Let *H*_PO,bet_ be the number of PO pairs found in offspring samples of size *n*_O,1_ and parent samples of size *n*_P,2_. Under the assumption that the total number of PO pairs between the two populations is much greater than *H*_PO,bet_, the distribution is approximated by a binomial form (i.e., *H*_PO,bet_ ∼ Binom[*π*_PO,bet_, *n*_O,1_*n*_P,2_]). Thus, the theoretical expectation of *H*_PO,bet_ is expressed as follows:

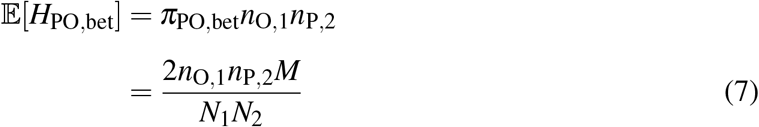

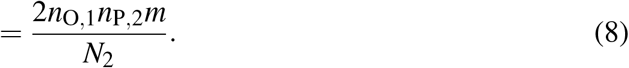

The observed number of PO pairs in a sample is defined by 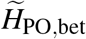, and 𝔼[*H*_PO,bet_] in Equations 7 and 8 are replaced by 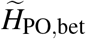, which generates the linear estimators of *M* and *m* as follows:

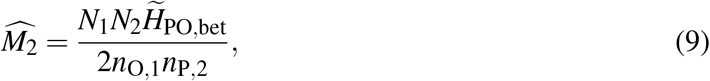

and

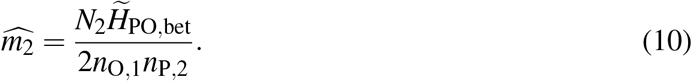

### 2.5 Required sample size

The proposed estimators are based on the observed number of kinship pairs. Their expected number is linearly determined by the number of pairwise comparison (Equations 2-3 and 7-8), which provides sample size guidance to ensure the condition that at least one or more kinship pairs can be found. The conditions are given as follows:

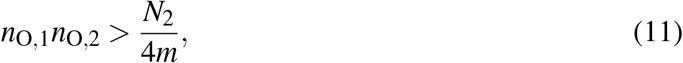

and

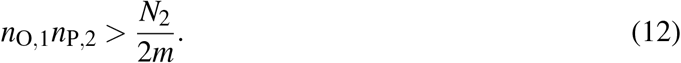

Generally, the required sample size to ensure the above conditions is 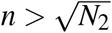, where *n* = *n*_O,1_ = *n*_O,2_ = *n*_P,2_. Note that a very small *m* value, which dramatically increases the required sample size, produces significant genetic differences between the two populations. In such cases, there are several methods to estimate migration rate using population genetics technique. Alternatively, we focus on moderate/large migration rate, which yields little genetic differences between the two populations.

Other guidance for the required sample size is obtained by an approximate lower bound on the coefficient of variation (CV) of *M*_1_ or *m*_1_, which is defined as 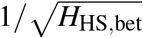. This is applied in the context of classic MR (Seber, 2002) or CKMR (Bravington, Skaug, & Anderson, 2016). For example, to achieve a 30% CV, the target of *H*_HS,bet_ is greater than 10, thereby providing the required sample size when the population parameters are given (e.g., *N*_1_, *N*_2_, and *M*). Similarly, the CV of *M*_2_ or *m*_2_, defined by 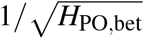, provides the sample size required to estimate *M*_2_ or *m*_2_.

### 2.6 Efficient use of kinship pairs found between populations

When both HS and PO pairs, which are found between populations, are available simultaneously, we can obtain efficient estimators by combined 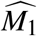 and 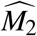 for migration number:

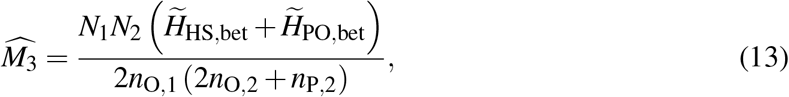

and by combining 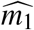 and 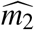 for the migration rate:

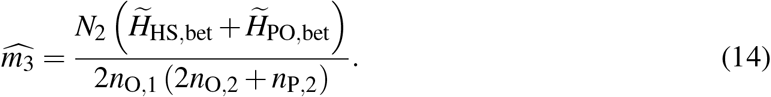

For those estimators, 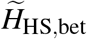 and 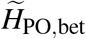 are weighted by 2*n*_O,2_ and *n*_P,2_, respectively.

### 2.7 Estimation of parent number by PO pairs in a population

To this point, we have stated that the formulation of the estimators, presented in Equations 4-5 and 9-10, is a function with parent numbers for each population (*N*_1_ and/or *N*_2_). In other words, such estimators are available only when the parent number or numbers are known. Here, we describe how we estimate unknown parent numbers by additionally using PO pairs to produce estimators for both migration number and rate that can be obtained from only the genetic data.

When PO pairs found in offspring samples and parent samples from population 2 in the second year are available, the standard parental number estimator can be obtained (Bravington, Skaug, & Anderson, 2016) as follows:

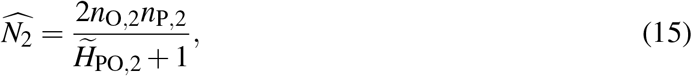

where 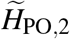 is the observed number of PO pairs found in the offspring samples of size *n*_O,2_ and parent samples of size *n*_P,2_. This term “+1” reduces bias, especially when 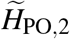 is small (e.g., Prystupa, McCracken, Perry, & Ruzzante, 2021), where a similar derivation of this bias correction is provided in the literature Akita (2020a). By replacing *N*_2_ by 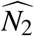 in Eqs. 13 and 14, we obtain the following estimators:

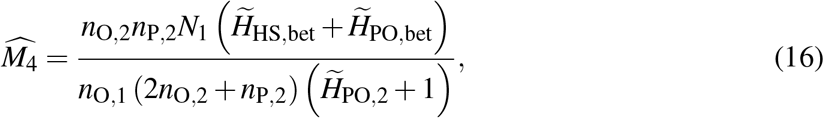

and

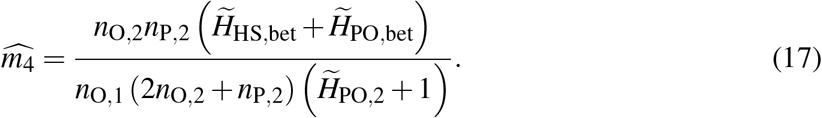

While the estimator of the migration number (Equation 16) requires the (unknown) parent number in population 1 (*N*_1_), the estimator of the migration rate (Equation 17) can be obtained using only the observed number of HS and PO pairs.

Similar to estimating *N*_2_, when PO pairs found in offspring samples and parent samples from population 1 in the first year are available, we can obtain the estimator of *N*_1_ as follows:

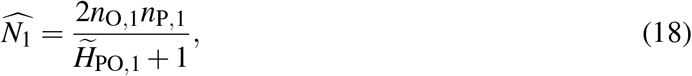

where 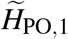 is the observed number of PO pairs in the offspring samples of size *n*_O,1_ and parent samples of size *n*_P,1_. By replacing *N*_1_ with 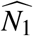 in Eq. 16, we obtain a migration number estimator that can be calculated from only genetic data, which is expressed as follows:

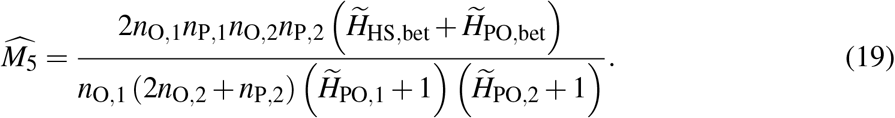

Table 2 summarizes the conditions for estimators in terms of whether *N*_1_ and/or *N*_2_ are known and which type of samples are required for estimation. In addition, Table 2 identifies the kinship type required to calculate the estimator.

### 2.8 Variation in reproductive potential among individuals and populations

The proposed estimators are derived under the assumption that reproductive potential is equal among individuals. As described in **Appendix 2**, this assumption can be relaxed, and the estimators still hold, which is exemplified in iteroparous species that may show significant variation in reproductive potential among individuals. In addition, the estimators hold for most cases where the mean reproductive potential differes between populations 1 and 2, e.g., when the environmental condition for reproductive success varies between populations. Here, the required condition is that the migration event does not depend on the degree of reproductive potential (refer to **Appendix 3** for additional details).

### 2.9 Individual-based model

We developed an individual-based model that tracks kinship relationships to evaluate the performance of the estimators. Here, the population structure was assumed to be identical to that in the development of the estimators. Population 1 and 2 comprised *N*_1_ and *N*_2_ parents with an equal sex ratio, and their offspring number was assumed to follow the geometric distribution with mean 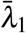 and 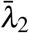 (i.e., Poisson reproduction with mean *λ*_*i*,1_ and *λ* _*j*,2_, which follows the exponential distribution with mean 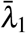 and 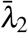), respectively. Migrant parents were selected randomly from population 1 at the end of the first year, and each offspring retained IDs of the parents, thereby making it possible to trace a HS and PO relationship.

Here, the parameter set (*N*_1_, *N*_2_, *M*, 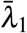, 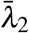, *n*_O,1_, *n*_O,2_, *n*_P,1_, *n*_P,2_) is given. We simulated a population history and a sampling process, which generates proposed estimators. Note that this process was repeated 1000 times, which allows us to construct the distribution of the estimators for each parameter set. All scripts (written by C++ and R languages) and documentation for these analyses are available at https://github.com/teTUNAakita/CKMRmig.

## 3 Results

We evaluated the performance of 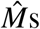 or 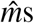 numerically for a case with variable reproductive potential among parents. Here, the scaled statistical properties of 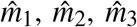 and 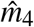 were completely the same as 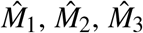, and 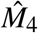; thus, hereafter, we only demonstrate the results for 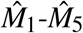. **Figure 2** shows the distribution of the relative bias of 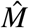 for limiting cases where parent and offspring sample numbers are identical (i.e., *n* = *n*_P,1_ = *n*_P,2_ = *n*_O,1_ = *n*_O,2_) and parent numbers in the two populations are also identical (i.e., *N* = *N*_1_ = *N*_2_). The relative bias is calculated by applying the outputs of the individual-based model, which is defined as “(averaged estimator − true value)/true value.” Refer to **Table S1** in **Supporting Information** for a full list of the parameter sets used to evaluate performance (relative bias and CV of 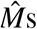).

**FIGURE 2.**
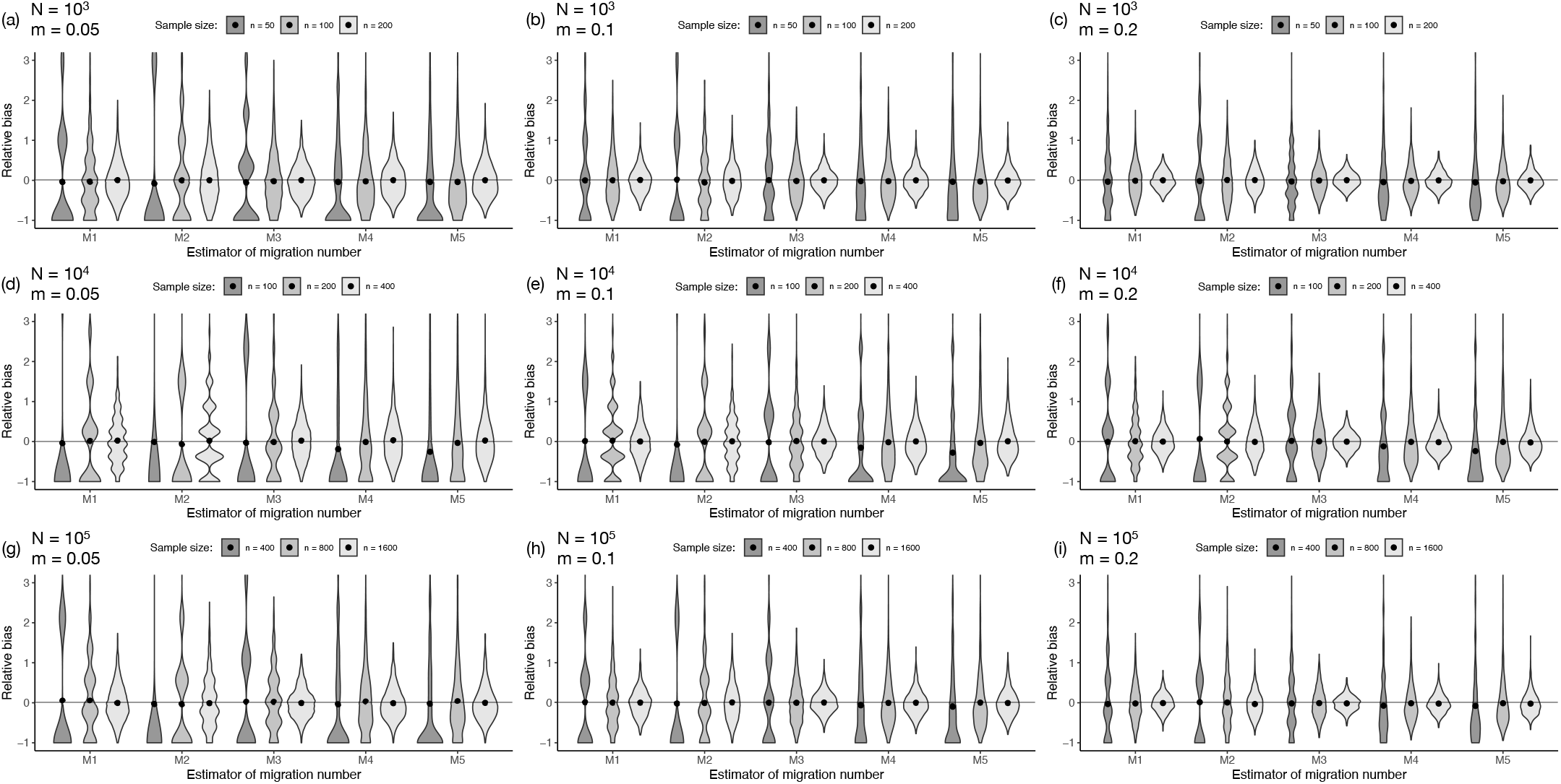
Violin plots showing the distribution of relative bias in the estimator of *M* for various sample sizes, parent numbers, and migration rates. Filled circles represent mean values. The sample sizes for parents and offspring are identical (i.e., *n* = *n*_P,1_ = *n*_P,2_ = *n*_O,1_ = *n*_O,2_), and the parent sizes in the two populations are identical (i.e., *N* = *N*_1_ = *N*_2_), as indicated in the legend. Migration rate *m* is specified by *N*_1_*/M*. For demonstration purposes, the upper side of the distribution is truncated, although the mean values are calculated (including the truncated values).

First, we evaluated the accuracy of 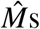 based on the relative bias. As expected, for most of the investigated parameter sets, we observed that their relative bias was less than 5%, as shown in **Table S1** in **Supporting Information**. Thus, when the assumptions that (i) offspring and parents are randomly sampled, (ii) kinships are detected without any error, and (iii) the migration event does not depend on the degree of reproductive potential, are satisfied, it is reasonable to consider 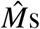 nearly unbiased estimators.

Next, we evaluated the precision of 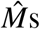 based on the CV value. **Table S1** in **Supporting Information** shows the CV value, and the violin plot in **Figure 2** visualizes the degree of precision. For each estimator 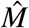, we found that precision increases with increasing sample size. Note that the total sample size depends on the estimator. For example, *n* = 50 in Fig. 2, the total sample sizes of 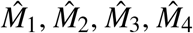, and 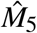 were 100 (= *n*_O,1_ + *n*_O,2_), 100 (= *n*_O,1_ + *n*_P,2_), 150 (= *n*_O,1_ + *n*_O,2_ + *n*_P,2_), 150 (= *n*_O,1_ + *n*_O,2_ + *n*_P,2_), and 200 (= *n*_O,1_ + *n*_O,2_ + *n*_P,1_ + *n*_P,2_), respectively.

As the sample size or number of migrants increase, precision increases, and the shape of the distribution asymptotically becomes symmetric (**Fig. 2**) because an increasing sample size or increasing number of migrants is likely to increase the observed number of kinship pairs found between the two populations (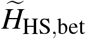 or 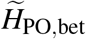) and decreases the variance of those kinship pair numbers. In addition, the number of PO pairs in a given population (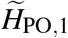 or 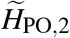) contributes to the precision of estimating *N*_1_ or *N*_2_, respectively, thereby providing 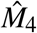 and 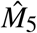 with a relatively high precision.

Note that our simulation can handle case with and without invasive sampling. Invasive sampling potentially affects the level of *m* because the sampled parents from population 1 have no chance to move to population 2; thus, such parents cannot be sampled. In this case, *m* might be defined by *M/*(*N*_1_ − *n*_*P*,1_) rather than *M/N*_1_, although this does not affect the estimator 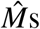, as demonstrated in **Table S1** (**Supporting Information**).

Finally, we investigated the case where the parental number in population 1 is much greater in population 2 (i.e., *N*_1_ ≫ *N*_2_); thus, migrants represent a large proportion of *N*_2_. For such cases, e.g., (*N*_1_, *N*_2_, *M*) = (10^4^, 10^3^, 500), we confirmed the similar property of 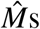 to the *N*_1_ = *N*_2_ situation, as shown in **Table S1** (**Supporting Information**), which suggests the robustness of 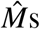 in the presence of heterogeneous population sizes.

## 4 Discussion

In this study, we theoretically developed estimators for the contemporary migration number 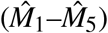 and migration rate 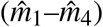 of parents between two predefined populations in iteroparous species. The proposed estimators are based on the known PO relationship and HS relationships observed between and within the two populations without any error in terms of kinship assignment. Users can select the appropriate estimator for cases where the parental number of population 1 (*N*_1_) and/or that of population 2 (*N*_2_) is known (**Table 2**). The performance of the estimator (accuracy and precision) was evaluated quantitatively by performing an individual-based simulation (**Fig. 2** and **Table S1** in **Supporting Information**). The proposed modeling framework utilizes several types of reproductive variations (i.e., the number of survived offspring per parent), including the variance of reproductive potential within and between populations, in consideration of several situations, including a body-size structure or environmental heterogeneity for reproductive success.

Our primary contributions are summarized as follows. First, we formulated the probabilities of kinship pairs randomly selected between two populations (*π*_HS,bet_ and *π*_PO,bet_), which provides migration number or rate estimators. While similar derivations may be found in the (nongenetic) MR method, these are limited to cases where sampling is non-invasive and adult individuals must be sampled. The proposed method can avoid these limitations, where such an advantages is characterized by the CKMR method for estimating population sizes. Second, we have demonstrated that the probabilities of kinship pairs are approximately independent of the reproductive potential; thus, information about it is not required to realize the estimation provided that migration is assumed to occur independently of the reproductive potential. This is a useful property of the estimators because it is natural that heterogeneity of reproductive potential within and between populations exits in iteroparous species but its observation is generally difficult. Third, we have demonstrated that estimators can be obtained using only genetic data (i.e., 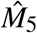 and 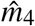). There are several advantages in using the proposed estimator rather than estimating *π*_HS,bet_ (and/or *π*_PO,bet_) separately (via a genetic method) and *N*_1_ (and/or *N*_2_) (via a nongenetic method), including simplified sampling processes and analyzing designs and the availability of a unified framework of genetic analyses for detecting HS and PO pairs (similar discussion is found for estimating the ratio of effective breeding size to the census size, *N*_b_*/N*, in Akita (2020b)).

To estimate the contemporary migration number or migration rate, our simulation-based results provide sample size guidance to ensure the required accuracy and precision, especially if the order of the number of migrant parents and parental sizes are approximately known (**Table S1** in **Supporting Information**). For example, when *m* = 0.1 and *N*_1_ = *N*_2_ = 10^3^, sampling 10% and 20% of the parents and an equal number of offspring in both populations leads to 70% and 39% CV of 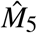, respectively (in the invasive sampling case). Even no information is available about these numbers, 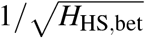 (or 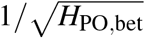) provides an approximate lower bound on the CV, which can be used as an indicator of the precision of 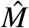 or 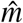 In addition, the condition that 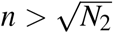 is also used as rule of thumb especially when planning a research project (*n* = *n*_O,1_ = *n*_O,2_ = *n*_P,2_; refer to Equations 11 and 12). Note that the guidance is to estimate the information about one-way migration between two populations; thus, estimating two-way migrations among two or three populations would require twice and three times the number of samples, respectively, compared to our sample size guidance.

We believe there are several scenarios where the proposed estimator may be beneficial. The first case is where there is a large number of migrants between populations, e.g., *M* > 100, which eliminates population differentiation and thus hampers the detection of the *M* via population genetics method if *M* is interpreted as an effective migration number. Information about movement between populations is essential to assess population dynamics in the conservation and management contexts, even if the migrant number is so large that it cannot be genetically assigned to two populations. The second case involves genetic monitoring conducted for each reproductive period (e.g., annually) because the proposed estimators consider information about contemporary migration that is explicitly specified the timing (e.g., year); thus, the time-series data for migration may reflect environmental changes, and this would provide insights into the underlying ecological processes. In addition, HS pairs found within the same cohort in the genetic population provide *N*_b_ (Wang, 2009; Waples & Waples, 2011), which is also used to assess genetic health. The third is the case involves the development of integrated models that combine several data sources into a single analysis (Maunder & Punt, 2013). An example of this direction is to provide additional information about migration to ongoing CKMR projects for stock assessment in fisheries (e.g., Bravington, Grewe, & Davies, 2016; Conn et al., 2020; Hillary et al., 2018; Prystupa et al., 2021; Trenkel et al., 2022), which would utilize kinship pairs for simultaneous estimation of population parameters. If the population structure is defined hypothetically, the data accumulated in such projects could be readily connected to our theory via likelihood methods. The fourth case is where sampling adults is difficult due to conservation practices or other reasons but instead sampling offspring is relatively easy. In such cases, although *N*_2_ (and *N*_1_) must be given externally to employ 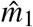 (and 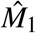), there is presently no existing method to estimate adult movement, especially when two populations cannot be genetically distinct, which indicates that the proposed method has the potential to expand the scope of current population monitoring techniques.

Finally, we discuss some caveats in terms of applying the proposed method. Our theory for developing the estimators assumes that kinships are detected without error. Many algorithms can be used to detect kinship pairs from single nucleotide polymorphisms or short tandem repeats (e.g., Huisman, 2017; Wang & Santure, 2009), although a HS pair requires many more DNA markers than a PO pair. In addition, iteroparous species may has several kinship types, e.g., half-uncle– nephew or half-cousins, which are expected to appear frequently and should be accurately differentiated from HS pairs. It is desirable to estimate in advance how many markers are required for kinship detection in the target populations associated with simulation of pedigree reconstruction (e.g., Anderson (in press)). The proposed estimators are limited to detecting parental movements on a period between breeding seasons in the given populations. Thus, the estimation of fine-scale spatiotemporal movements, which is available to integrate data sources (Thorson et al., 2021), is beyond the scope of this paper. Although population dynamics models with coarser spatial resolution than the spatial scale of environmental layers are frequently used in assessment models, i.e., the target application of the proposed method, the proposed estimators require pre-specification of the population structure. Kinship relationships with sample location information potentially realize the ability to explore a plausible population structure by estimating the migration number or rate between hypothetical populations, which is expected to contribute the determination of a management unit. In addition, the current theory behind the estimators does not assume desynchronized reproduction within a population (e.g., skip spawning), correlation between individual mobility and fertility, and nonrandom sampling of parents (e.g., mothers with a large number of offspring are likely to be sampled). These issues will be the focus of future work.

## ACKNOWLEDGMENTS

The author thanks N. Shinohara, N. Suzuki and Y. Tsukahara for their fruitful discussions. The author also thanks W.M. Iwasaki for helping with the C++ coding. This work was supported by JSPS KAKENHI Grant Numbers 19K06862 and 20H03012.

## AUTHOR CONTRIBUTIONS

**Tetsuya Akita:** Conceptualization (lead); formal analysis (lead); funding acquisition (lead); methodology (lead); writing–original draft (lead); writing–review and editing (lead).

## DATA ACCESSIBILITY

No datasets were generated or analyzed in this study.

## APPENDIX 1 Reproductive potential

Here, we introduce the concept of the reproductive potentials of parents *i* and *j* in the population 1 and 2, respectively, which are defined as the expected number of surviving offspring at the given sampling time, denoted by *λ*_*i*,1_ and *λ* _*j*,2_ (*i* = 1, …, *N*_1_ and *j* = 1, …, *N*_2_). Reproductive potential is determined by several factors, including the parent age, weight, and residence time in the spawning ground. Note that the magnitude of this parameter includes information about offspring survival rate, the number of days after egg hatching, and the number of eggs. This implies that the parameter reflects the sample timing. Also note that the modeling framework does not depend on whether the reproductive potential is heritable or not.

## APPENDIX 2 Derivation of Equations 1 and 6 when reproductive potential is variable among parents

In the main text, we ignore the variation of reproductive potential among parents (i.e., both *λ*_*i*,1_ and *λ* _*j*,2_ are constant) to derive π_HS,bet_ and π_PO,bet_. Here, let *k*_*i*,1_ and *k* _*j*,2_ be the number of surviving offspring of parents *i* and *j* at sampling time in populations 1 and 2, and assumed to follow a kind of discrete distribution (e.g., Poisson or negative binomial distribution) with mean *λ*_*i*,1_ and *λ* _*j*,2_, respectively. Without loss of generality, we set the index such that parents with *i* = 1 to *M* in population 1 and *j* = 1 to *M* in population 2 are identical migrants. For example, the parent with *i* = 1 reproduces *k*_1,1_ offspring (in population 1) and then reproduce *k*_1,2_ offspring (in population 2) after migration. Given *k*_*i*,1_ and *k* _*j*,2_, the conditional probability that two offspring sampled in populations 1 and 2 share a HS relationship is expressed as follows:

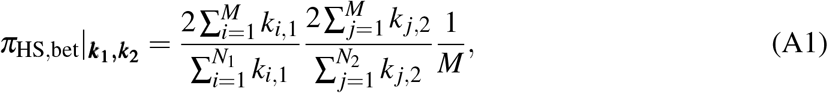

where 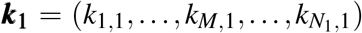 and 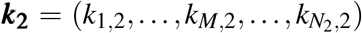. Note that *k*_*i*,1_ and *k* _*j*,1_ are a random variables with mean *λ*_*i*,1_ and *λ* _*j*,2_, respectively. By taking the expectation over the distribution of the offspring number, the conditional probability is given approximately as follows:

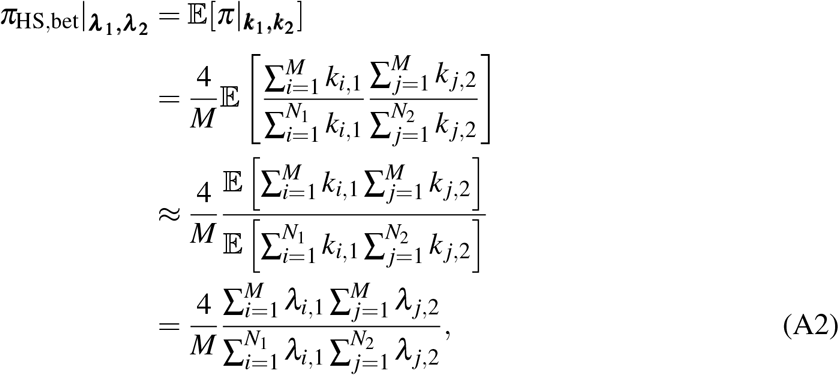

where 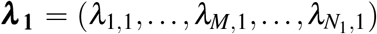 and 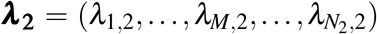. From the second to third lines, we use the approximation that 𝔼[*g*1(*k*)*/g*2(*k*)] ≈ 𝔼[*g*_1_(*k*)]*/* 𝔼[*g*_2_(*k*)]. From the third to forth lines, we use the relationship that 𝔼[*k*_*i*,1_*k* _*j*,2_|_*λi*,1,*λj*,2_] = 𝔼[*k*_*i*,1_|_*λi*,1_] 𝔼[*k*_*j*,2_|_*λj*,2_], which implies the statistical independence of the offspring number before and after parental movement. In other words, the conditional probability is not affected by 𝕍 [*k*|_*λ*_]. Here, we assume that *λ*_*i*,1_ and *λ* _*j*,2_ are also random variables that are followed by an arbitrary function with mean 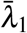 and 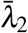, respectively. By taking the expectation over *λ* and applying a similar approximation, the unconditional probability is given as follows:

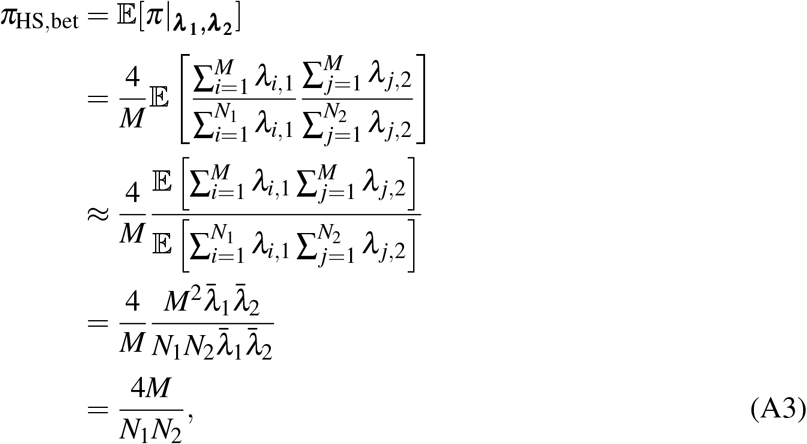

which provides the same formulation described in Equation 1. Note that, from the third to forth lines, we assume that *λ*_*l*,1_ and *λ*_*l*,2_ (*l* = 1, …, *M*) are independent variables (i.e., 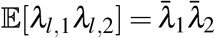), which implies variable reproductive potential of an identical parent before and after migration.

Next, we derive the probability that offspring sampled in population 1 and a parent sampled in population 2 share a PO relationship (*π*_PO,bet_) under the flexible settings of *λ*, which is similar to the derivation of *π*_HS,bet_. The conditional probability is expressed as follows:

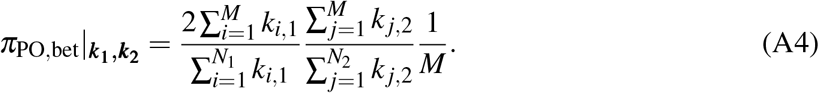

Here, by taking the expectation over *k* and *λ*, in the same manner as noted above, the unconditional probability is approximately given as follows:

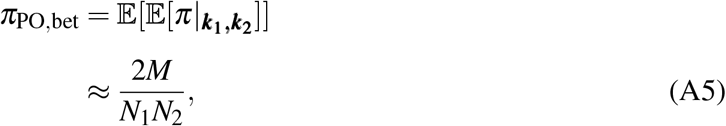

which provides the same formulation described in Eq.6.

## APPENDIX 3 Difference in reproductive potential between migrants and non-migrants

In the derivation of *π*_HS,bet_ and *π*_PO,bet_ in **Appendix 2**, we ignored covariation between migration and reproductive potential. Here, we consider the case where migrants have a distinguishable distribution of reproductive potential from nonmigrant parents. Let *λ*_*i*,1_, *λ* _*j*,2_, *λ*_*M,l*,1_ and *λ*_*M,l*,2_ be the reproductive potential of nonmigrant parents in population 1 and 2 and migrants in the populations 1 and 2 with mean 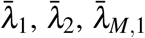, and 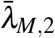, respectively. Under this setting, *π*_HS,bet_ can be expressed as follows:

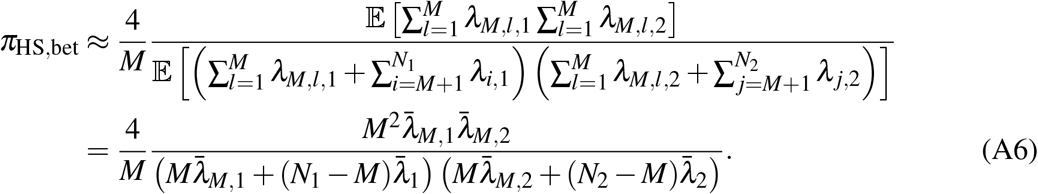

If 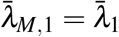 and 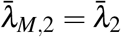, Equation A6 can be reduced Equation A3. This formulation includes reproductive potential terms such that the distinct reproductive potential between migrants and nonmigrants eliminates the usefulness of the proposed HS-based estimators (this is also applied to the proposed PO-based estimators).

